# Alterations in DNA 5-hydroxymethylation Patterns in the Hippocampus of an Experimental Model of Refractory Epilepsy

**DOI:** 10.1101/2023.10.03.560698

**Authors:** Rudhab Bahabry, Rebecca M. Hauser, Richard G. Sánchez, Silvienne Sint Jago, Lara Ianov, Remy J. Stuckey, R. Ryley Parrish, Lawrence Ver Hoef, Farah D. Lubin

## Abstract

Temporal lobe epilepsy (TLE) is a type of focal epilepsy characterized by spontaneous recurrent seizures originating from the hippocampus. The epigenetic reprogramming hypothesis of epileptogenesis suggests that the development of TLE is associated with alterations in gene transcription changes resulting in a hyperexcitable network in TLE. DNA 5-methylcytosine (5-mC) is an epigenetic mechanism that has been associated with chronic epilepsy. However, the contribution of 5-hydroxymethylcytosine (5-hmC), a product of 5-mC demethylation by the Ten-Eleven Translocation (TET) family proteins in chronic TLE is poorly understood. 5-hmC is abundant in the brain and acts as a stable epigenetic mark altering gene expression through several mechanisms. Here, we found that the levels of bulk DNA 5-hmC but not 5-mC were significantly reduced in the hippocampus of human TLE patients and in the kainic acid (KA) TLE rat model. Using 5-hmC hMeDIP-sequencing, we characterized 5-hmC distribution across the genome and found bidirectional regulation of 5-hmC at intergenic regions within gene bodies. We found that hypohydroxymethylated 5-hmC intergenic regions were associated with several epilepsy-related genes, including *Gal*, *SV2,* and *Kcnj11* and hyperdroxymethylation 5-hmC intergenic regions were associated with *Gad65*, *TLR4*, and *Bdnf* gene expression. Mechanistically, *Tet1* knockdown in the hippocampus was sufficient to decrease 5-hmC levels and increase seizure susceptibility following KA administration. In contrast, *Tet1* overexpression in the hippocampus resulted in increased 5-hmC levels associated with improved seizure resiliency in response to KA. These findings suggest an important role for 5-hmC as an epigenetic regulator of epilepsy that can be manipulated to influence seizure outcomes.

## Introduction

Temporal lobe epilepsy (TLE) is a neurological disorder characterized by recurrent unprovoked seizures arising from the temporal lobe, frequently in the hippocampus. Epigenetic mechanisms play a crucial role in the regulation of the epileptic state [35]. Particularly, DNA cytosine methylation is significantly altered in several models of epilepsy in both the brain and blood [4, 6, 10, 22, 57, 74, 75, 85, 86, 88, 94, 96, 105, 109, 118, 131]. However, DNA methylation patterns differ by the epilepsy model used, showing that seizure origins are crucial in the consideration of epileptic DNA methylation patterns [22]. The interpretation of DNA methylation changes in epilepsy is challenged by the main use of the traditional bisulfite sequencing method, which does not differentiate between the two main forms of methylated DNA in the brain, 5-methylcytosine (5-mC) and 5-hydroxymethylcytosine (5-hmC) [43]. In this study, we address for the first time the contribution of 5-hmC to the epileptic methylome.

DNA can be methylated through the activity of DNA methyl transferases (DNMTs) at cytosine residues, and methylated DNA (5-mC) typically represses transcription of the associated gene [8, 40, 91]. 5-mC can be oxidized by TET (Ten-Eleven-Translocation) enzymes to form DNA hydroxymethylation (5-hmC) [46, 117]. This oxidation both acts as an intermediate step in active DNA demethylation and as a stable hydroxymethyl mark added to the DNA [3, 36, 126]. In contrast to 5-mC, which is typically associated with gene repression, 5-hmC is mostly associated with gene activation. 5-hmC is enriched at transcriptionally active sites such as gene bodies and poised and active enhancers [112, 115]. DNA 5-hmC can act by preventing repressive proteins with methyl binding domains (MBDs) from binding to and repressing gene expression, therefore increasing gene expression [49]. 5-hmC can also act as a *cis*-element regulating gene expression through the binding of transcription factors (TFs) to 5-hmC at gene regulatory regions. In addition, 5-hmC is negatively correlated with H3K27me3-marked and H3K9me3-marked repressive genomic regions suggesting that 5-hmC may contribute to gene expression in a histone dependent manner [33, 68, 119]. In addition, 5-hmC at exon/intron junctions is thought to play a role in gene splicing [54].

5-hmC is more abundant in the brain than in other tissue types by approximately ten-fold, while 5-mC is more consistently enriched throughout tissue types [26, 60]. It is thought that active DNA methylation/demethylation is necessary for the regulation of neuronal activity. Previous studies have shown that 5-hmC is crucial for neuronal maturation and function, with 5-hmC levels increasing up to ten-fold over the course of development and enriched at genes with synapse-related functions [33, 116]. 5-hmC and TET enzymes have a crucial role in normal brain functions such as memory and memory acquisition [7, 12, 28, 31, 50, 63, 96, 104, 125]. In addition, 5-hmC also has been associated with numerous neurological disorders, including Rett syndrome, Alzheimer’s disease, autism, and Huntington’s disease [14, 47, 62, 83, 95, 113, 124, 133, 135]. Consequently, the methyl-binding protein MeCP2 is abundant in the brain and dysregulated in Rett syndrome. MeCP2 acts to regulate gene expression through interactions with methylated DNA [83]. Also interestingly, MeCP2 binds to both 5-mC and 5-hmC, and MeCP2 binding appears to prevent conversion of 5-mC to 5-hmC [77, 83]. This suggests unique roles for methyl binding proteins in genome and epigenome regulation dependent upon underlying DNA methylation type.

Due to 5-hmC’s abundance in the brain and its role in regulating neuronal activity, it is crucial to consider the contribution of 5-hmC when interpreting previously reported bisulfite sequencing data. In this study, we sought to map 5-hmC across the epileptic epigenome using 5-hydroxymethylcytosine methylated DNA immunoprecipitation sequencing (hMeDIP-seq). We observe significant genome-wide loss of 5-hmC with no significant changes to global 5-mC levels hippocampal tissue samples from TLE patients and from TLE mice. However, this trend is not reflective of the 5-hmC at the gene-specific level. The majority of gene-associated 5-hmC loss or gain occurs within gene bodies. Moreover, we provide evidence that DNA hydroxymethylation level changes can impact acute seizures. The results from this study take a closer look at DNA hydroxymethylation in epilepsy and further support the importance of DNA 5-hmC in the brain and as a critical epigenetic regulator in epilepsy worthy of further studies.

## Materials and methods

### Animals and Epilepsy Model

Male Sprague-Dawley rats received from Harlan weighing 125-150g at the time of arrival were used for these experiments. Animals were housed in pairs in plastic cages and had access to water and NIH-31 lab rat diet ad libitum with a 12:12hr light/dark cycle. All procedures were approved by the University of Alabama at Birmingham Institutional Animal Care and Use Committee and done in accordance with the National Institute of Health and ethical guidelines. Animals were handled by investigators for one week after arrival. Half of the animals were intraperitoneally (IP) injected with saline (vehicle), and the other half with 10mg/kg of kainic acid (KA) (0222, Tocris, Minneapolis, MN, USA). Behavioral seizure severity was measured using the Racine scale. 1: mouth and face clonus and head nodding; 2: clonic jerks of one forelimb; 3: bilateral forelimb clonus; 4: forelimb clonus and rearing; 5: forelimb clonus with rearing and falling [79, 101]. The onset of status epilepticus (SE) was defined as the time from KA injection to the start of continuous seizure activity with scores of 4 or 5 in the Racine scale. The animals providing samples for mass spectrometry analysis were sacrificed at 8 weeks post KA. The cohort used for hydroxymethylated DNA immunoprecipitation (hMeDIP-seq) was administered IP injections of saline once a day for two weeks after two months post KA. Animals were euthanized by rapid decapitation 24 hours following the last injection, and their hippocampi were removed and subdissected in ice-cold oxygenated artificial cerebral spinal fluid (ACSF). The cornu ammonis (CA) region was collected and frozen on dry ice. Mass spectrometry analysis was performed as previously described [50].

### KA challenge test

The animals used for *Tet1* manipulation experiments from the different treatment groups received challenging doses of KA 10 mg/kg IP injected at 30-minute intervals until SE was reached. Animals’ seizure behavior was observed and scored on the Racine scale as described above. Animals were euthanized by rapid decapitation 1 hour following SE for the *Tet1* KD experiment and 4 hours following the *Tet1* overexpression experiment.

#### Stereotactic delivery of siRNA and lentivirus

Rats were anesthetized with isoflurane (induction:5%, maintenance:2%) and placed in a stereotaxic frame. The hair on the scalp was shaved and the skull was exposed and leveled by placing bregma and lambda in the same horizontal plane, then a small craniotomy was performed. For experiments involving siRNA delivery, Accell SMARTpool siRNAs (Dharmacon) targeting Tet1 (#E-094677-00-0005) or negative control (#D-001910-10-05) into the dorsal hippocampus using stereotaxic coordinates (AP: −5.6 mm, ML: ± 4.5 mm, DV: −5.00 mm) relative to bregma. For experiments involving lentivirus delivery, plasmids were obtained from addgene a full-length human TET1 gene with a Flag and HA tag (#49792) and scrambled TALE-TET1CD (#49944) for negative control. Both genes: Full-size TET1 and TALE-TET1CD, were sequentially cloned into appropriate LV cloning vectors at UAB Neuroscience NINDS Vector and Virus Core C. Injections into the dorsal hippocampus using stereotaxic coordinates (AP: −5.6 mm, ML: ± 4.5 mm, DV: −5.00 mm) relative to Bregma. The infusion was given over 10 minutes (0.1 μl/minute) for a total volume of 1 μl per side. After each injection, the needle was left in place for an additional 10 minutes to allow for diffusion of the siRNA and was then slowly removed. The incision was closed using sutures and Vetbond.

### Hydroxymethylated DNA immunoprecipitation (hMeDIP) and Library Prep

Dissected CA samples were homogenized in Tris/EDTA (TE) buffer made from 1% 1M Tris pH 8.1 and 0.2% containing 1% SDS and 100µg of proteinase K (EO492, ThermoFisher Scientific) and incubated for 2hrs at 55°C. DNA was extracted through phenol/chloroform/isoamyl alcohol and ethanol precipitation. 400ng of DNA were used for hMeDIP assays. Samples were sheared to ∼300bp using 25 cycles on a Bioruptor (Bioruptor XL, Diagenode, Lome, AT) at high power, denatured at 95°C for 15 minutes, and diluted with IP buffer (16mM Tris, 1.2mM EDTA, 167mM NaCl, 1% SDS, 1% Triton). DNA was then added to MagnaChip magnetic protein A/G beads (16-663, Millipore, Burlington, USA) overnight that were incubated with either 5-hmC, or IgG the day before (2µg of anti –IgG (ab171870, Abcam) and 2µg of 5-hmC (39791, Active Motif, Carlsbad, USA)). The DNA bead complexes were washed with low salt buffer (0.1% SDS, 1% Triton X-100, 0.4% 0.5M EDTA, 2% 1M Tris-HCl pH 8.1, 0.875% w/v NaCl in DDH2O), high salt buffer (0.1% SDS, 1% Triton X-100, 0.4% 0.5M EDTA, 2% 1M Tris-HCl pH 8.1, 2.74% w/v NaCl in DDH2O), LiCl buffer (1.06% w/v LiCl, 0.1% NP-40, 1% Sodium Deoxycholate, 0.2% 0.5M EDTA, 1% 1M Tris-HCl pH8.1 in DDH2O), and TE buffer. Samples were then extracted with TE buffer containing 1% SDS and 100µg of proteinase K for 2hrs at 55°C and 10 minutes at 95°C. DNA was extracted and purified by phenol/chloroform/isoamyl alcohol and then ethanol precipitation. Library preps of the immunoprecipitated DNA samples were made using the ThruPLEX DNA-seq Kit (R400427, Rubicon Genomics, Ann Arbor, MI, USA), and sequencing was performed at UAB’s Heflin Genomics core at ∼15 million single end 75bp reads.

### hMeDIP-seq bioinformatics

The FASTQ files were uploaded to the UAB High-Performance Computer cluster for bioinformatics analysis with the following pipeline built in the Snakemake workflow system (v5.2.2): first, quality and control of the reads were assessed using FastQC, and trimming of the bases with quality scores of less than 20 and adapters were performed with Trim_Galore! (v0.4.5). Following trimming, the reads were aligned with Bowtie2 [64](v2.3.4.2, with option ‘--very-sensitive-local’ set) to the University of California Santa Cruz (UCSC) rat genome (rn6), which resulted in an average mapping rate of 96.20%. BAM files were sorted and indexed with SAMtools(v1.6) and logs of reports were summarized and visualized using MultiQC (v1.6). BAM files were imported into a local RStudio session (R version: 3.5.0) for the identification of differentially methylated regions (DMRs) with MEDIPS (v.1.34.0)[72] across two

DMR criteria: 1) the entire promoter (defined as 5000 bp upstream of the gene’s transcriptional start site (TSS) and the entire gene-body 2) genome-wide quantification using windows of 500 bp for visualization purposes of regions outside of gene-bodies and promoters). The ‘BSgenome.Rnorvegicus.UCSC.rn6’ (v.1.4.1) library was used as the reference. For the quantification of DMRs across promoters and gene-bodies, reference ‘bed’ files (from UCSC) of each region were implemented as the region of interest (ROI) in MEDIPS. Thus, MEDIPS datasets were created with the following parameters to the ‘MEDIPS.createROIset’ function: ROI=<promoter or gene-body references>, extend=0, shift=0, uniq= 1e^-3^ (to remove duplicates), paired=FALSE.

Computation of differential methylation was executed with the ‘MEDIPS.meth’ function using the following parameters: p.adj=”BH” (Benjamini-Hochberg method for p-value adjustment), diff.method=“edgeR”, MeDIP=TRUE, CNV=FALSE, minRowSum=10, diffnorm=“quantile”, CSet=CS (created with ‘MEDIPS.couplingVector(pattern = “CG”)’ with one of the control samples provided to ‘refObj‘), and MSet1 represented the ES group while MSet2 represented the CS group. Promoters and gene-bodies were called to be statistically significant if they contained an adjusted p-value < 0.05 and minimum mean count of 10 in one of the two groups. To generate genome-wide DMRs, MEDIPS datasets were created with the following parameters: extend = 300, shift = 0, uniq = 1e^-3^, window size = 500 and paired = FALSE. Differential methylation was computed in the same manner as promoters and gene bodies.

The circos plot was generated by Circos (v0.69-5) [61] using the rn6 ideogram obtained from the UCSC genome browser. The genome profile plots across the genic (gene-body), promoters, intergenic and CpG islands regions were generated with deeptools (v3.5.1) [102]. The input data for the profile plots were bigwig files generated from the genome-wide quantification analysis with a coverage cutoff of minimum mean count of 10 in one of the two groups (chromosome M was excluded). No statistical cutoff was applied for the generation of genome profile plots. The reference for intergenic region was created with BEDtools (v2.26.0) with ‘bedtools complement’ to generate a bed file of regions across the genome without genic and promoter regions. The CpG islands reference was acquired from UCSC, and region references were provided to deeptools’ ‘computeMatrix’ function via the −R parameter. Non-default arguments included ‘--skipZeros -bs 50’. For the CpG islands plot, upstream and downstream areas were extended by 4000bp. CpG shores and shelves were labeled as up to 2000 bp and 2000-4000bp up- and downstream of CGI respectively.

Gene ontology information was collected using WebGestalt[71]. Genes inputted were genes with an adjusted p-value of ≤ 0.05 and a log2FC of at least |0.75|. Seizure and epilepsy associated genes were taken from DisGeNET RDF v7.0 [98] using search terms “seizure” and “epilepsy.” For candidate genes an adjusted p-value ≤ 0.05 and a log2FC of at least |0.9| was used excluding chrM. Volcano plots were created using EnhancedVolcano (v1.12.0).

Average cell type gene expression data was collected using the Allen Brain Map whole cortex & hippocampus – 10X genomes (2020) with 10X – smart-seq taxonomy (2021) scRNA-seq dataset accessed November 3, 2021. The mean trimmed mean expression from cells and the metadata of class labels “Glutamatergic” and “GABAergic” were used to calculate the mean of GABAergic neuron and glutamatergic neuron gene expression. The mean trimmed mean from cells with the metadata subclass labels “Astro,” “Oligo,” and “Endo” were used to calculate the mean of gene expression in astrocytes, oligodendrocytes, and endothelial cells[128]. Graphical figures were created with BioRender.com. GraphPad Prism version 9.2.0 was used to create bar charts and heat maps. Gene maps were created using IGV version 2.11.2.

### Quantitative RT-PCR

Real-time PCR amplification of CA cDNA was performed on the Biorad CFX-96 Real-time system using TaqMan® Fast Advanced Master Mix and TaqMan® Gene expression assay using the following protocol: UNG activation at 50.0°C for 2 min, then polymerase activation at 95.0°C for 20 s, denature at 95.0°C for 3 s, followed by an Anneal/Extend at 60°C for 40 cycles. *Hprt1* expression was used to normalize gene expression. Cycle threshold (Ct) values were analyzed using comparative Ct method to calculate differences in gene expression between samples. Primers are as specified: *Hprt1* assay ID: Rn01527840_m1 VIC-MGB, *Sv2a* assay ID: Rn00589491_m1 FAM-MBG, *GAL* assay ID: Rn00583681_m1 FAM-MBG,

*Kcnj11* assay ID: Rn01764077_s1 FAM-MBG, *BDNF* assay ID: Rn02531967_s1 FAM-MBG, *TLR4* assay ID: Rn00569848_m1 FAM-MBG, *GAD2* assay ID: Rn00561244_m1 FAM-MBG.

### Human resected samples

Human samples were obtained from patients with medically intractable epilepsy undergoing elective neurosurgical resection of an epileptogenic hippocampus. All patients gave their informed consent, before surgery, for the use of the resected brain tissue for scientific studies. Human samples used for mass spectrometry were provided by Kristen O. Riley, MD from the Department of Neurological Surgery and G Yancy Gillespie, MD from the Wallace Tumor Institute at UAB, and from Tore Eid M.D. from Yale School of Medicine. Rosalinda Roberts, Ph.D. provided non-epileptic post-mortem samples. Patient demographics and pharmacological history were previously described in supplementary table 2 [106].

### Statistical analyses

Data is presented as a mean and standard error of the mean (SEM) where appropriate. Experiments were analyzed using unpaired t-tests performed by GraphPad Prism version 9.2.0. Adjusted p-values for sequencing data were calculated using edgeR [13]. Gene ontology FDRs were obtained from WebGestalt [71].

## Results

### 5-hmC but not 5-mC is lost with epilepsy

To determine if 5-mC or 5-hmC changes globally in the epileptic hippocampus, we investigated the total 5-hmC and 5-mC levels in both resected epileptic human and rodent hippocampus through mass spectrometry (**Figure 1c, f)**. Analysis of epileptic human tissue showed no overall changes in levels of DNA 5-mC **(Figure 1d)**, yet it revealed a global loss of 5-hmC **(Figure 1e)**. To determine if the change in DNA hydroxymethylation existed in rodent models of epilepsy, rats were injected intraperitoneally with KA to induce status epilepticus. Eight weeks following status epilepticus the animals were fully epileptic, and the Cornu Ammonis (CA) region of the hippocampus was collected and analyzed for DNA methylation using mass spectrometry **(Figure 1f)**. Mirroring the human data, we observed no change in DNA 5-mC levels **(Figure 1g)** and a decrease in total levels of DNA 5-hmC **(Figure 1h)**

**Figure 1.**
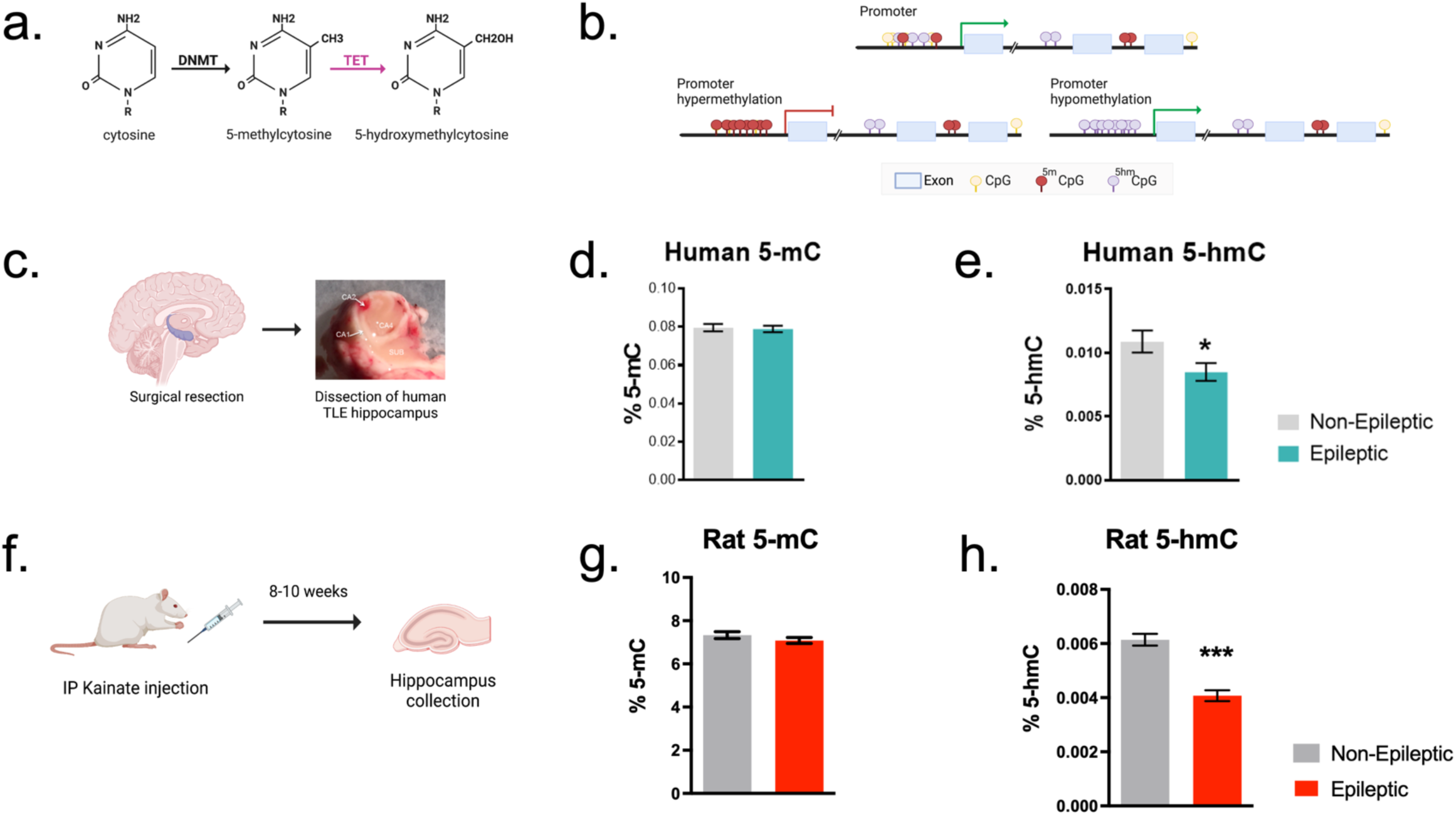
Total 5-hmC but not 5-mC is Lost with Epilepsy a) DNMTs methylate cytosine residues on DNA which can then be hydroxymethylated by TETs. b) Illustration of the different ways that DNA methylation and hydroxymethylation regulate gene expression. c) Human hippocampi were collected from epilepsy patients undergoing surgical resection of their hippocampus d) Mass spectrometry analysis showing the total percentage of DNA 5-mC in epilepsy and e) 5-hmC percentage from resected epileptic human hippocampus and post-mortem controls. f) Timeline of epilepsy rat model. Kainic acid was IP injected into rats to induce status epilepticus. After 8-10 weeks, the animals develop spontaneous seizures and the CA region of the hippocampus is collected. g) Mass spectrometry analysis total DNA 5-mC and h) 5-hmC percentage from epileptic rat hippocampus. *p<0.05, **p<0.01, ***p<0.001 unpaired t-test.

### Epileptic loss of 5-hmC occurs primarily at intergenic regions

While our mass spec data showed global changes in 5-hmC, we wanted to determine the genomic distribution of 5-hmC in epileptic animals, the CA region of the hippocampus in control and epileptic rats was used for 5-hmC meDIP-sequencing. The genome was split into 500bp windows, of which were 5517 hypermethylated and 6485 hypomethylated. Agreeing with our mass spectrometry data, epileptic rats had less DNA 5-hmC overall levels. Excluding mitochondrial DNA, 290, 2455, and 2772 windows were hypermethylated in gene promoters, bodies, and intergenic regions, respectively. 300, 2567, and 3618 windows were hypomethylated in gene promoters, gene bodies and intergenic regions, respectively. The intergenic windows, annotated gene bodies, and annotated gene promoters were then mapped across the genome. **(Figure 2)**. When examining only differentially hydroxymethylated (DhM) gene bodies, no general trend towards loss or gain of 5-hmC is noticeable **(Figure 2).** Similarly, gene promoter regions also show no clear bias towards loss or gain of 5-hmC except for chromosome X, which shows no hyper-hydroxymethylated promoters **(Figure 2).** Interestingly, the majority of 5-hmC loss is seen within intergenic regions **(Figure 2).**

**Figure 2.**
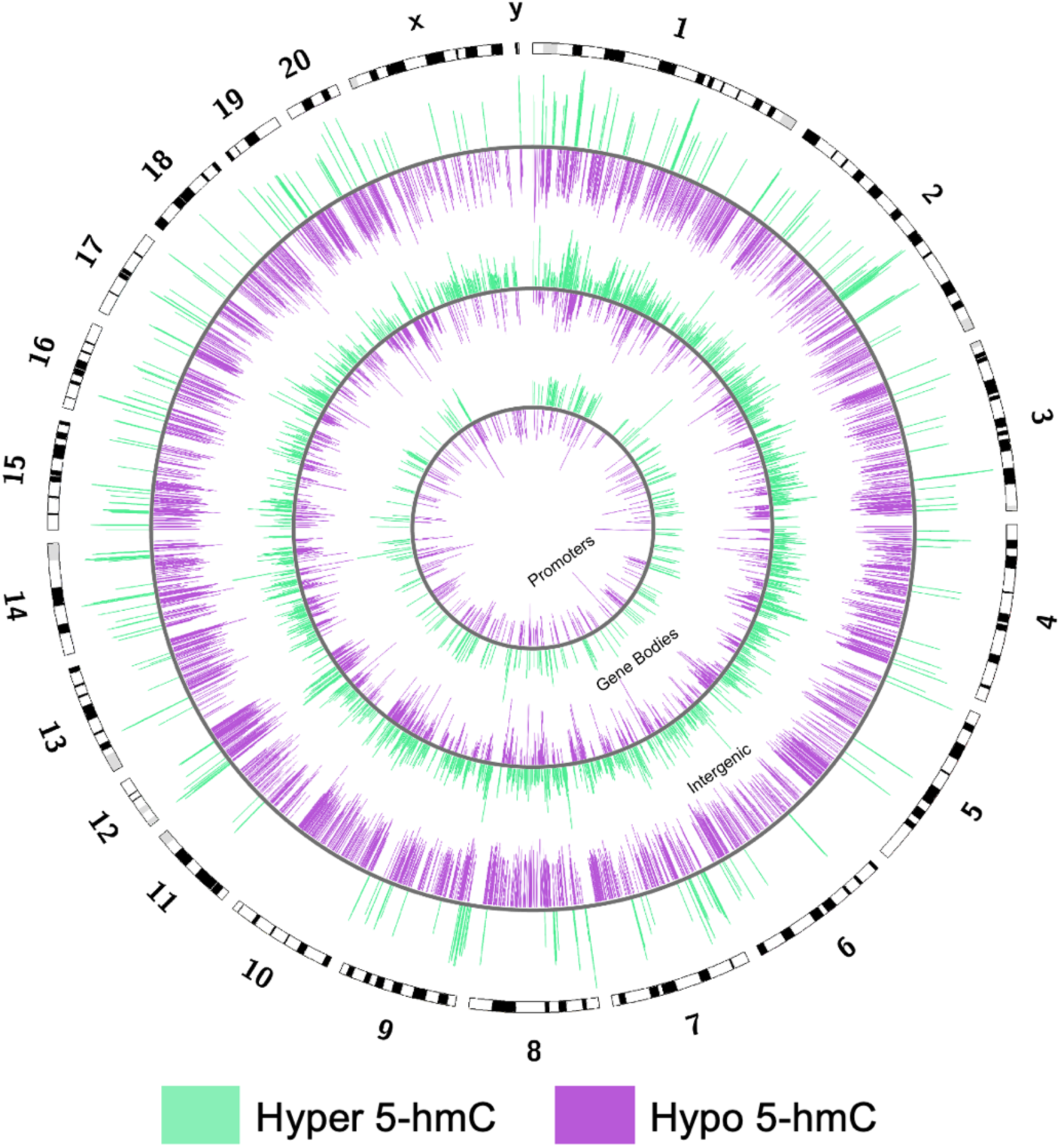
Differential 5-hmC Windows Across Genome Regions Circos plots of gene promoter and gene body regions of interest and intergenic 500bp windows 5-hmC showing Log2FC of DhMRs from epileptic rats relative to non-epileptic animals. Purple indicates regions with reduced 5-hmC relative to non-epileptic animals and green regions with increased 5-hmC.

### Epilepsy alters 5-hmC DNA methylation at gene bodies and gene promoters

To identify genes regulated by 5-hmC in epilepsy, we looked specifically at gene bodies and gene promoters with significant increases or decreases of 5-hmC. The majority of gene-associated DNA 5-hmC changes were seen at the gene bodies, with 2531 genes identified as hypomethylated and 2455 genes hypermethylated in epileptic rats relative to non-epileptic controls **(Figure 3a)**. In addition, we found 298 hypomethylated gene promoters and 290 hypermethylated promoters in epileptic rats **(Figure 3b)**. We also looked at 5-hmC in selected genes involved in neuronal and synaptic regulation and found altered Hydroxymethylation levels in their gene bodies and promoters **(Supplementary Material, Fig. S1a,b)**. Relative to CpG islands (CGI), the majority of differentially hydroxymethylated regions (DHMRs) were found at CpG shores, agreeing with past CpG methylation studies **(Figure 3c)** [44, 87], epileptic rats showed less 5-hmC than control animals with enrichment at CGI regions and a relatively even distribution of 5-hmC across CpG shores and shelves **(Figure 3g).** Across the average promoter, gene body, and intergenic region, there is a consistent decrease in 5-hmC occupancy in epileptic rats **(Figure 3d-f)**.

**Figure 3.**
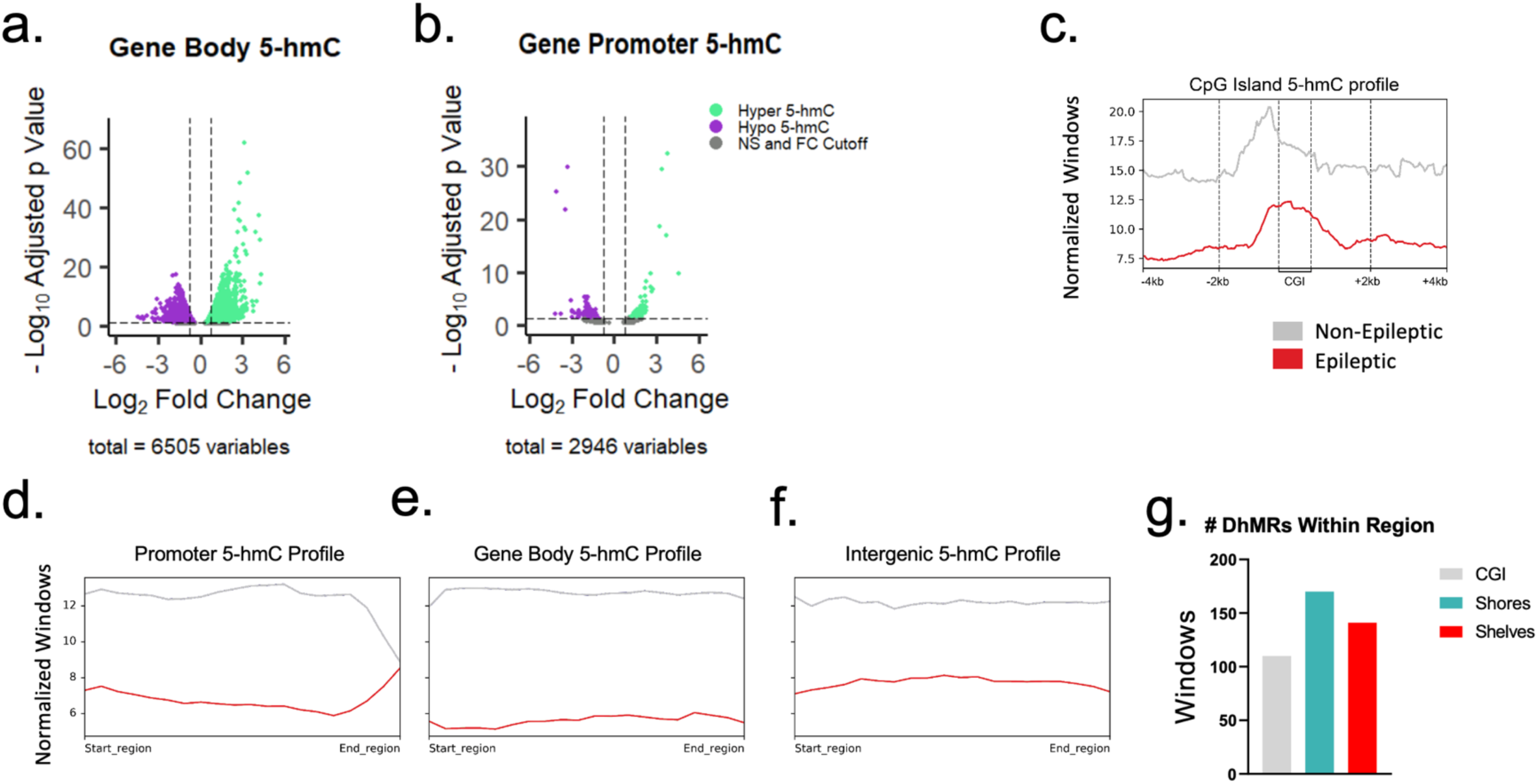
5-hmC Distribution a) Volcano plot showing hypo (purple) and hyper (green) hydroxymethylated gene bodies and b) Gene promoters. Adjusted p-value cutoff of 0.05 and fold change cutoff of Log2FC 0.75 are indicated by dotted lines. Genes with an adjusted p-value of more than 0.05 or a Log2FC 0.75 less than 0.75 are indicated in gray. c) Profile plots of average 5-hmC distribution across gene promoters d) gene bodies, e) intergenic regions, and f) CpG islands. g) Total number of differentially methylated 500bp windows of epileptic relative to control rats located within CGI (CpG Islands), CGI shores, and CGI shelves.

### Differentially hydroxymethylated genes are involved in several epilepsy-associated pathways

To address what pathways differentially methylated genes are involved in, the most significantly hypo- and hypermethylated gene bodies with a log2 fold change absolute value of at least 0.75 were submitted for gene ontology analysis (**Figure 4)**. We found that these genes were involved in several pathways, including GABA signaling as the top hit for hypermethylated gene bodies and several ion transport pathways in hypomethylated genes. GABA signaling is known to be important in epilepsy, with loss of GABA promoting seizure activity and increasing GABA having an antiseizure effect. In addition, multiple GABA receptor mutations result in clinical epilepsy [2, 38, 80, 99]. Ion channels are also critical for a variety of neuronal functions such as action potential propagation, resting membrane potential, and intercellular signaling and mutations in several channel genes cause genetic epilepsy [9, 66, 93]. KEGG pathway analysis showed enrichment of genes involved in retinol metabolism and nicotine addiction in hypermethylated genes. Retinoic acid acts as an anti-seizure therapeutic, and nicotine can be a pro-convulsant but has anti-convulsant effects in treating certain epilepsies [23, 103, 127]. Hypomethylated genes were associated with several signaling pathways such as sodium reabsorption regulated by aldosterone, JAK-STAT signaling, and cAMP signaling. Loss of sodium causing hyponatremia can result in seizures, and hyperaldosteronism can result in seizures [89, 130]. Both the JAK-STAT and cAMP signaling pathways are implicated in seizures [30, 34, 78, 82, 84]. Associated Reactome pathways of hypermethylated genes include GABA-A receptor activation and Retinoic Acid synthesis and pathways, and hypomethylated genes were enriched in PIP3 activation of AKT signaling and intracellular signaling. The PIP3 pathway has also been associated with epilepsy progression with seizures, causing a reduction in PIP3 [11]. Overall, enrichment analysis by GO terms, KEGG pathway, and Reactome pathways show overlapping patterns with DhMR enrichment in genes associated with GABA receptors, retinoic acid metabolism, and intracellular signaling. Furthermore, there are genes in multiple molecular pathways that are impacted by changes in DNA hydroxymethylation such as Neurogenesis, Synaptic signaling, microRNA, Learning & memory and Genes regulated by nuclear calcium signaling **(Supplementary Material, Fig. S2)**. This indicates that 5-hmC might be involved in the regulation of multiple important epilepsy-associated pathways and that 5-hmC could serve as a critical regulator of epilepsy development.

**Figure 4.**
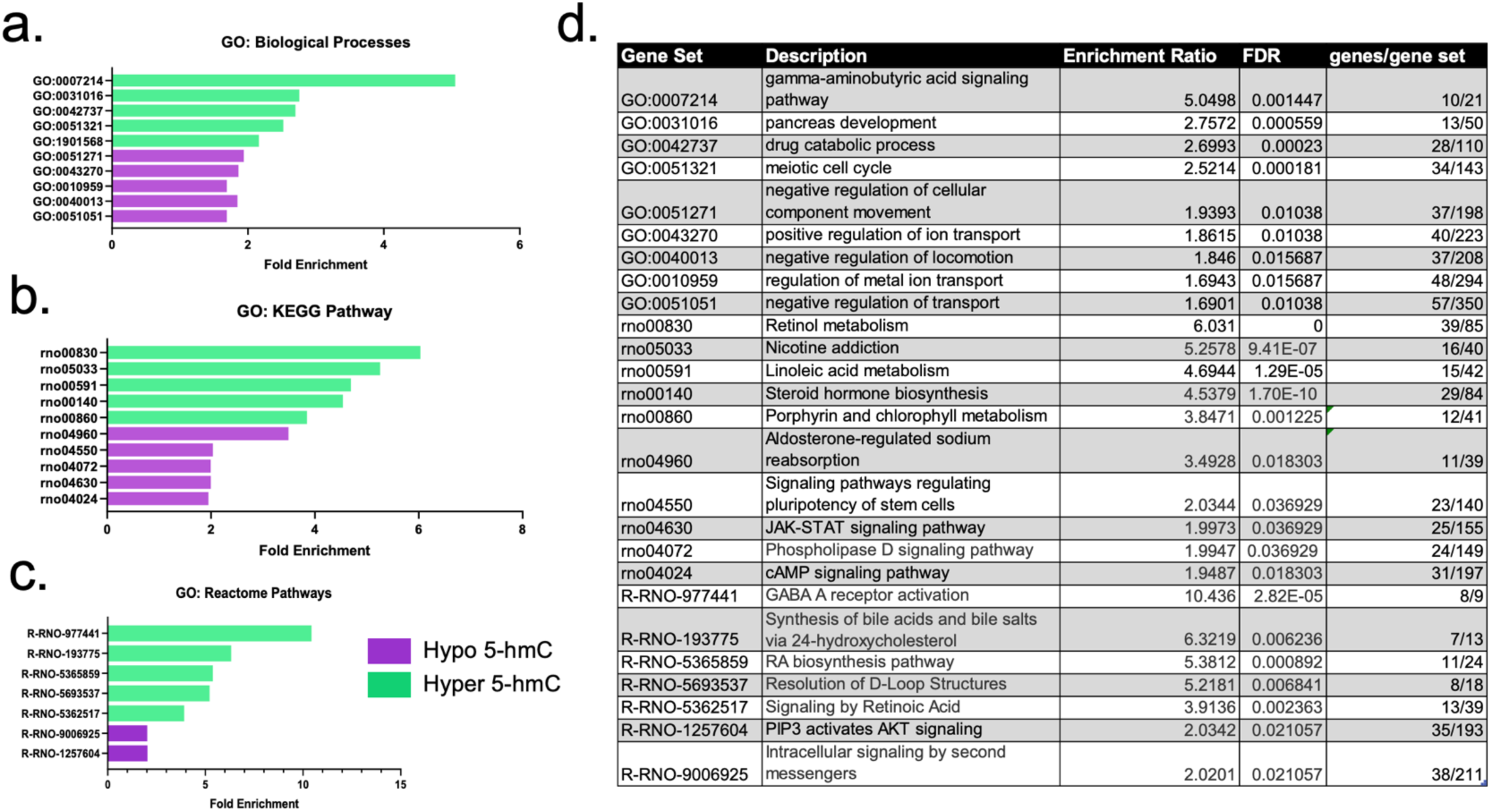
Pathway Enrichment of Differentially Hydroxymethylated Gene Bodies a) Fold enrichment of gene ontology pathways for hyper (green) and hypo (purple) hydroxymethylated genes for KEGG pathways, b) Biological Processes, c) and Reactome Pathways for gene bodies with an adjusted p-value <0.05 and a Log2FC of at least 0.75. d) Table describing gene set ids, descriptions, enrichment ratios, and FDR of gene ontology graphs in a, b, and c.

### DhMRs affect genes in a variety of cell types

To develop a list of candidate genes for 5-hmC regulation in epilepsy, we filtered our list of differentially methylated genes for genes known to be associated with epilepsy or seizures. From this, 69 genes were identified. This was further narrowed by the genes with the largest fold change in hydroxymethylation compared to non-epileptic animals to generate a list of 26 candidate genes **(Supplementary Material, Fig. 3a, b, c)**. Multiple cell types are involved in the hippocampal pathology of TLE. Seizures result from an overabundance of excitatory glutamatergic transmission and a loss of GABAergic inhibition. In epilepsy with hippocampal sclerosis, there is gliosis of the hippocampus resulting in the increase of hippocampal astrocytes. Astrocytes regulate glutamate in the brain as part of the tripartite synapse and the glutamate-glutamine cycle through glutamine uptake and conversion to glutamate [39]. As a crucial component of the myelin sheath, oligodendrocytes are essential regulators of neuronal transmission. In epilepsy, myelination abnormalities and damaged myelin sheaths have been reported [45, 73, 108, 129, 134]. Endothelial cells line blood vessels and play a vital role in maintaining the blood-brain barrier, which is disrupted in epilepsy [29]. Using the Allen Brain Institute’s scRNA-seq cell type database [128], we compared our list of candidate genes to the average expression in the following cell types: astrocytes, glutamatergic neurons, GABAergic neurons, oligodendrocytes, and endothelial cells. Of the top 26 hits, 7 genes are highly expressed in glutamatergic neurons, 6 in GABAergic neurons, 1 in astrocytes, 1 in endothelial cells, and 13 genes had no to low average expression in the cell types examined **(Supplementary Material, Fig. 3d).**

### Epileptic gene regulation by DhMRs

To examine if epileptic 5-hmC resulted in functional changes, we chose 3 hypo- and hypermethylated genes from our candidate genes to measure gene expression via qPCR: *Sv2a, Gal, Kcnj11* **(Figure 5)***, Bdnf, Tlr4*, and *Gad2 (***Figure 6)**. *Sv2a* encodes synaptic vesicle protein 2a, which is the binding site for the antiepileptic drug levetiracetam. Galanin (G*al*) is an endogenous neuropeptide that acts to inhibit seizures [32, 67]. Mutations in the potassium channel subunit Kir6.2 (K*cnj11*) are known to result in genetic epilepsy [27, 123]. Brain derived neurotrophic factor (*Bdnf*) has antiseizure effects, however, infusion of BDNF can also lead to seizures [24, 45, 65, 92, 107]. Toll-like receptor 4 (*Tlr4*) activation increases inflammation which can lead to seizures [1, 20, 37, 55, 122]. Glutamate decarboxylase 2 (*Gad2*) converts glutamate into GABA, and loss of expression and activity of GAD2 is associated with seizure activity [18, 51, 132]. Typically, 5-hmC correlates with an increase in gene expression. All 3 hypomethylated genes had reduced gene expression in the hippocampus **(Figure 5a-i)**. Interestingly, all the hypermethylated genes tested also showed a significant reduction in gene expression **(Figure 6a-i)**. This could potentially be explained by the heterogeneity of cell types in the tissue sent for meDIP-seq, regulation of these genes through other epigenetic mechanisms, or involvement of intergenic 5-hmC regulation.

**Figure 5.**
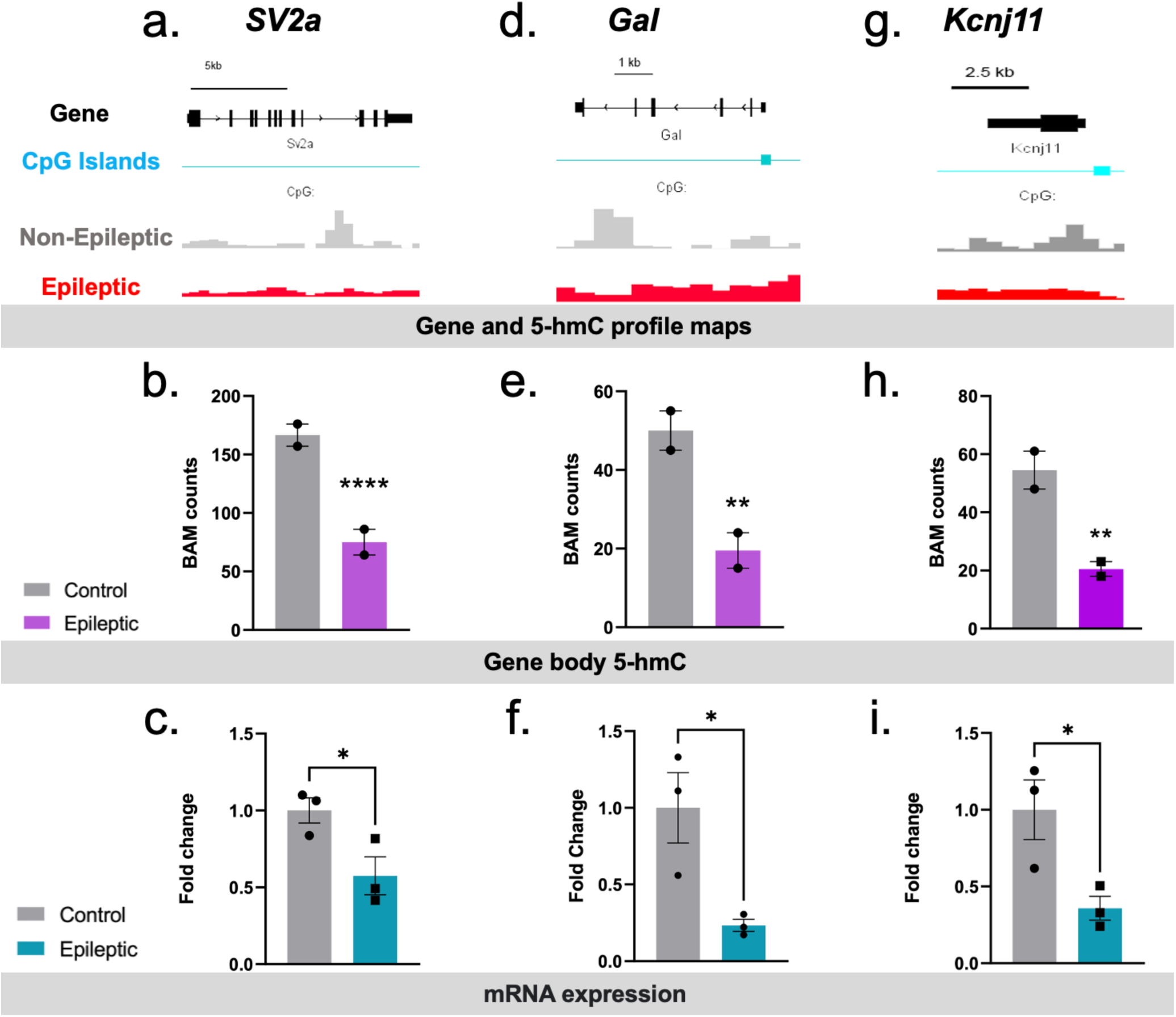
Expression of Hypohydroxymethylated Genes a) Gene profile map of *SV2a*. Non-epileptic 5-hmC profiles are shown in gray, epileptic profiles are red. CpG islands are shown in blue, and the gene body exons and introns are shown in black. b) BAM counts of *SV2a* 5-hmC in epileptic (purple) and nonepileptic (gray) animals. c) qPCR gene expression of *SV2a* in epileptic (blue) and nonepileptic (gray) animals. d) Average gene profile of *Gal* 5-hmC in epileptic and nonepileptic animals. e) BAM counts of *Gal* 5-hmC in epileptic and non-epileptic rats. f) *Gal* mRNA levels in epileptic rats. g) Profile of 5-hmC distribution at the *Kcnj11* gene. h) BAM counts of *Kcnj11* 5-hmC in epileptic and non-epileptic animals. i) *Kcnj11* mRNA levels of in epileptic rats. *p<0.05, **p<0.01, ***p<0.001, ****p<0.0001 unpaired t-test (mRNA expression graphs), edgeR adjusted p-value (BAM counts).

**Figure 6.**
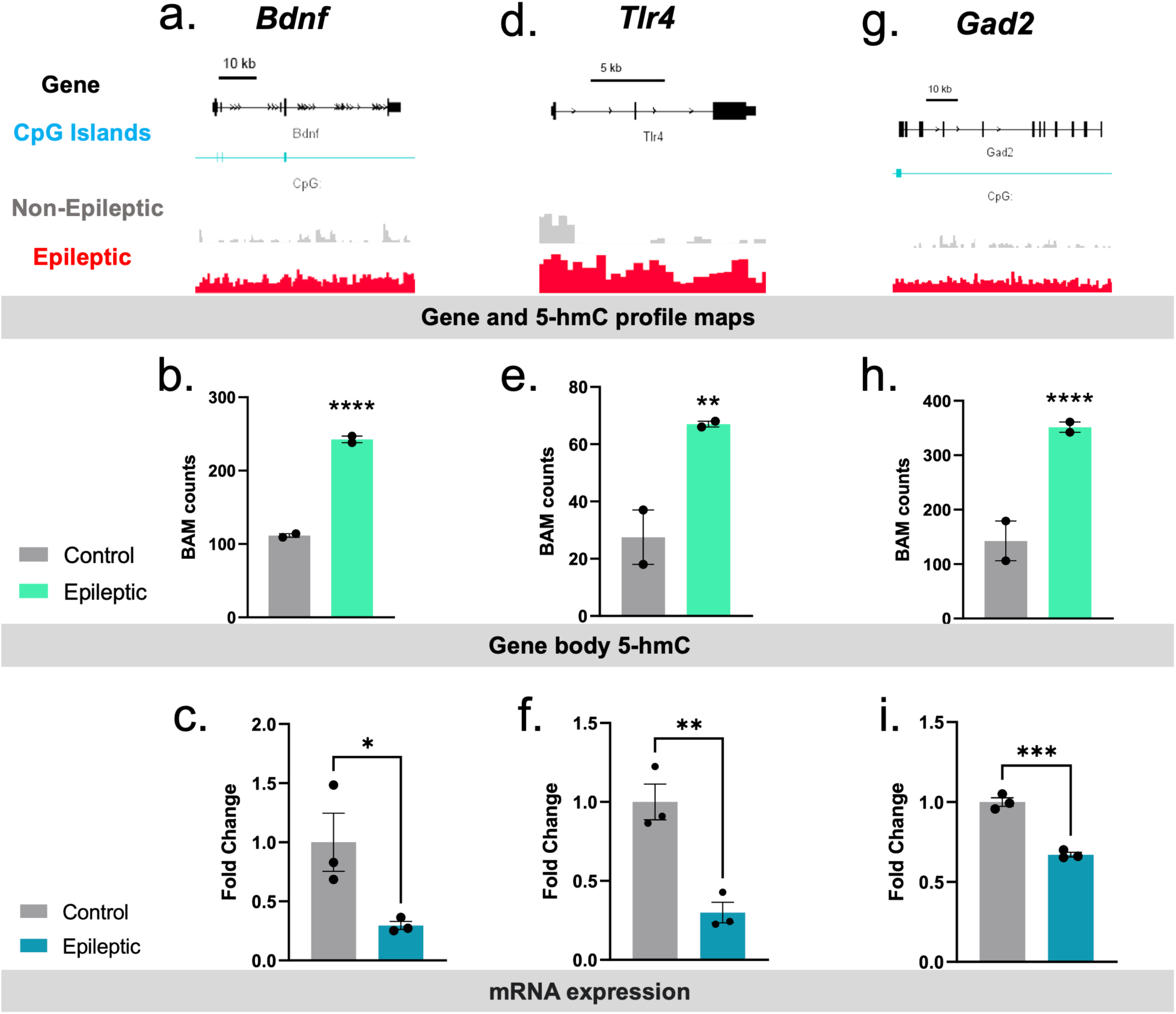
Expression of Hyperhydroxymethylated Genes a) Gene profile map of *Bdnf*. Non-epileptic 5-hmC profiles are shown in gray, epileptic profiles are red. CpG islands are shown in blue, and the gene body exons and introns are shown in black. b) BAM counts of *Bdnf* 5-hmC in epileptic (green) and nonepileptic (gray) animals. c) qPCR *Bdnf* gene expression in epileptic (blue) and nonepileptic (gray) animals. d) Average gene profile of *Tlr4* 5-hmC in epileptic and nonepileptic animals. e) BAM counts of *Tlr4* 5-hmC in epileptic and non-epileptic rats. f) *Tlr4* mRNA levels of in epileptic rats. g) Profile of 5-hmC distribution at the *Gad2* gene. h) BAM counts of *Gad2* 5-hmC in epileptic and non-epileptic animals. i) *Gad2* mRNA levels in epileptic rats. *p<0.05, **p<0.01, ***p<0.001, ****p<0.0001 unpaired t-test (mRNA expression graphs), edgeR adjusted p-value (BAM counts).

### Knockdown of Tet1 increases seizure susceptibility

From this study and others, it’s been well established that there are significant changes in DNA methylation and hydroxymethylation occurring in the epileptic hippocampus affecting different biological processes and gene expression [21, 56, 97, 111]. So, we hypothesized that reduction of *Tet1* in the dorsal hippocampus will reduce 5-hmC levels and influence seizure sensitivity. To do that, we used *in vivo* siRNA-mediated knockdown of *Tet1* in the dorsal hippocampus of Sprague Dawley rats and after 5 days, we induced seizures acutely by IP KA injections. Rats were observed for behavioral seizures and scored according to the Racine scale **(Figure 7a)**. We witnessed that animals treated with *Tet1* siRNA had faster latency to forelimb clonus following the KA challenge compared to the ones infused with scrambled siRNA **(Figure 7d)**. However, no difference was detected in the latency to SE between the two groups **(Figure 7e)**. ELISA analysis of hippocampal tissues revealed no change in 5-mC DNA methylation levels during SE **(Figure 7b)**. In contrast, there was a significant reduction in 5-hmC levels in *Tet1 siRNA-injected* rats compared to the scrambled siRNA-treated controls **(Figure 7c)**. Here, we illustrated the direct relationship between DNA hydroxymethylation and seizures, since we found that *in vivo* knockdown of *Tet1* in the dorsal hippocampus leads to the reduction of 5-hmC levels which subsequently increased seizure vulnerability following KA challenge.

**Figure 7.**
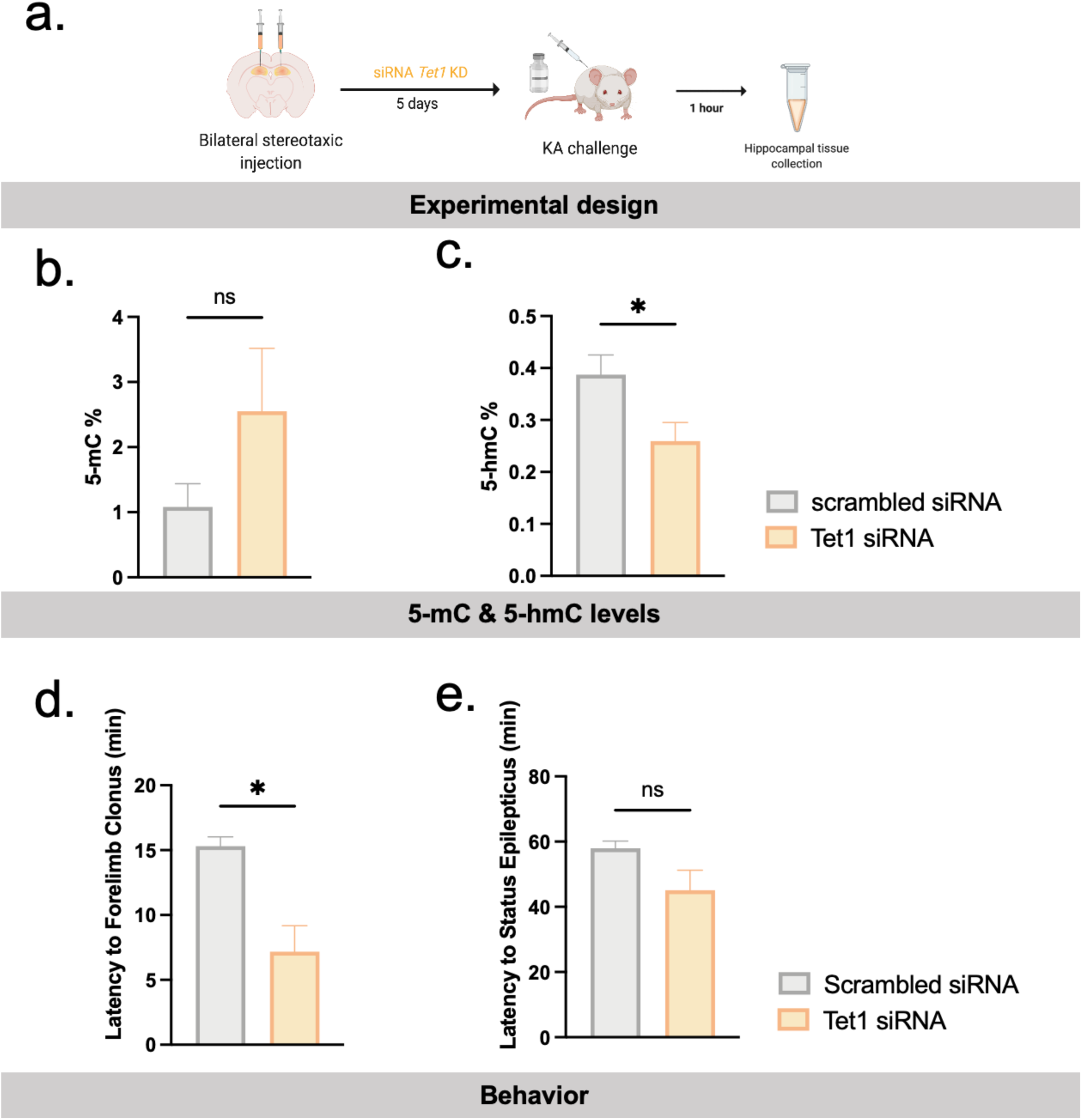
*Tet1* Knockdown is sufficient to increase seizure susceptibility a) Tet1 knockdown experimental timeline schematic. b) Latency to forelimb clonus, c) Status epilepticus and d) Duration of severe seizure in minutes after scrambled siRNA or Tet1 siRNA. e) ELISA analysis of hippocampal tissues of 5-mC and f) 5-hmC DNA methylation levels. n=3-5, *p<0.05.

### Overexpression of Tet1 promotes resilience to seizure

Next, we examined if increasing DNA hydroxymethylation will improve the seizure phenotype. So, we used *in vivo* Lentivirus-mediated overexpression of *Tet1* in Sprague Dawley rats delivered to the dorsal hippocampus and after 3 weeks, we induced seizures acutely by IP KA injections. Rats were observed for behavioral seizures and scored according to the Racine scale **(Figure 8a)**. No difference was detected in the latency to forelimb clonus in both groups, *Tet*1 overexpression and inactive *Tet1* rats **(Figure 8d)**. A significant delay was detected in the latency to SE observed in *Tet1* overexpression treated group compared to the inactive *Tet1* treated control rats **(Figure 8e)**. ELISA analysis of hippocampal tissues revealed no change in 5-mC DNA methylation levels during SE **(Figure 8b)**. In contrast, there was a significant increase in 5-hmC levels in *Tet1* overexpression injected rats compared to the inactive *Tet1* treated control rats **(Figure 8c)**. Our results show that the increase in DNA hydroxymethylation associated with the overexpression of *Tet1 in vivo* provided resilience to seizure by extending the time window to reach full SE following the KA challenge. Altogether, these behavioral and molecular changes suggest alterations of *Tet1* in the dorsal hippocampus play a critical role in seizure vulnerability and resilience.

**Figure 8.**
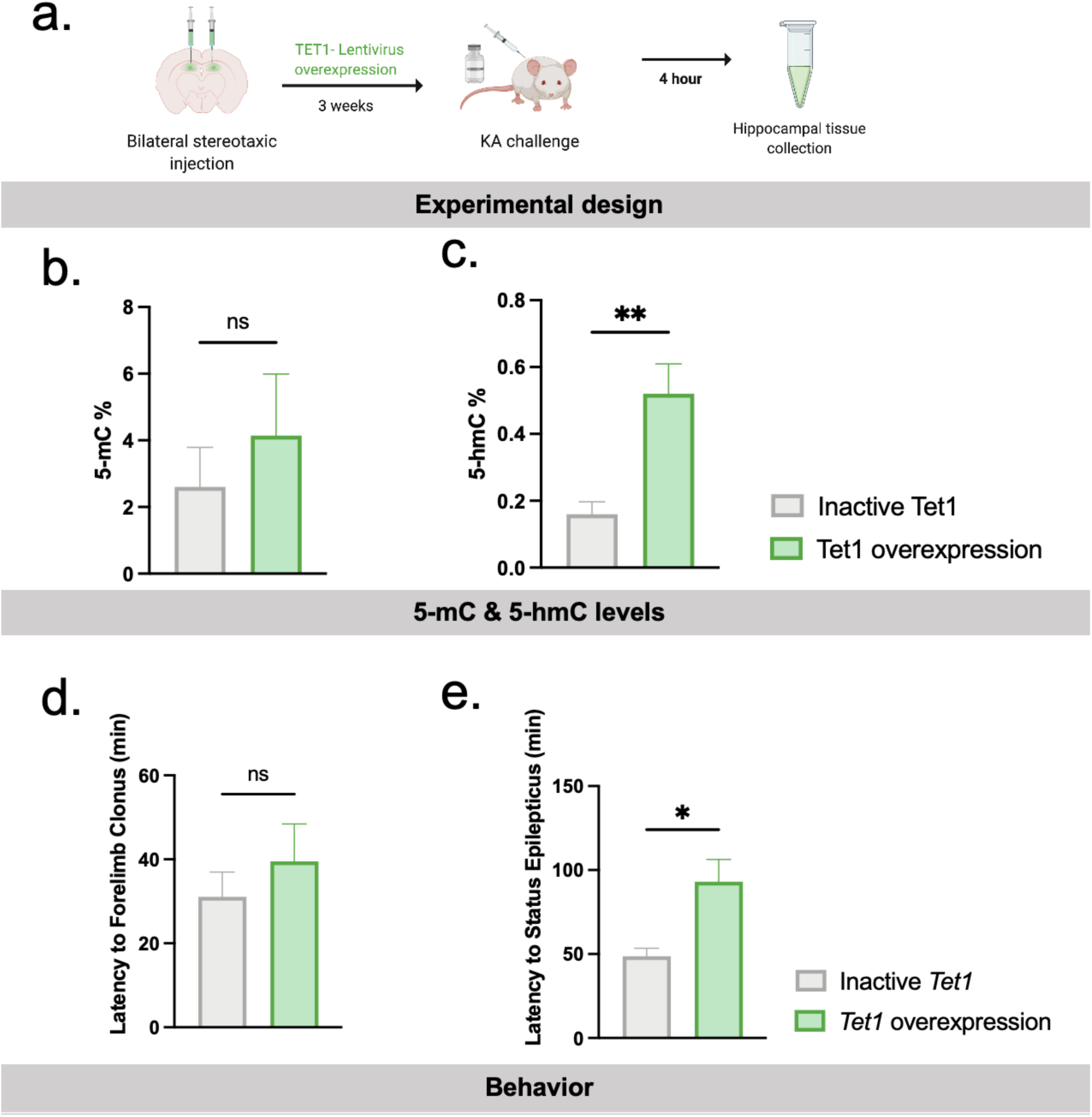
*Tet1* Overexpression delays status epilepticus onset and promotes seizure resiliency a) Tet1 overexpression experimental timeline schematic. b) Latency to forelimb clonus, c) Status epilepticus and d) Duration of severe seizure in minutes after inactive Tet1 or Tet1 overexpression. e) ELISA analysis of hippocampal tissues of 5-mC and f) 5-hmC DNA methylation levels. n=4-6, *p<0.05.

## Discussion

Expression of TET enzymes have been found to be altered in human TLE and rodent TLE models and following status epilepticus, a prolonged seizure event [19, 50, 105, 111]. However, very little is known about 5-hmC distribution pattern across the epigenome in TLE. In this study, we performed DNA 5-hmC hMeDIP-sequencing to establish a 5-hmC genome map and showed that overall DNA methylation changes are largely driven by 5-hmC instead of 5-mC in chronic epilepsy. We observed loss of DNA 5-hmC and no change in total 5-mC in human TLE hippocampus and the KA rat model of epilepsy. Total loss of 5-hmC is also described in TLE by de Nijs et al [19], the electrical kindling model of epilepsy by [111], and 24 hours following flurothyl-induced seizures [50], suggesting that DNA 5-hmC loss in the hippocampus might be a consistent hallmark of epilepsy pathology and that 5-hmC could be more important than 5-mC as a crucial epigenetic regulator of epilepsy.

We characterized the distribution of genomic 5-hmC in epilepsy and identified several DNA 5-hmC regulated candidate genes in epilepsy. Specifically, we examined the gene expression of three candidate genes with significantly hypomethylated gene bodies: *Sv2a, Gal,* and *Kcnj11*. These genes show decreased mRNA levels in the hippocampus of epileptic rats. We also examined the expression of three hypermethylated genes: *Bdnf, Tlr4,* and *Gad2*. All of the hypermethylated genes tested showed a significant loss in gene expression contrary to the expected increase with increased 5-hmC occupancy at the gene body. Interestingly, several studies have shown an increase in *Bdnf* gene and protein expression [24, 45, 96, 105]. *Bdnf* has several splice variants that are differentially expressed in response to neuronal activity such as with and epilepsy and memory formation [15, 76, 81, 82, 90, 96, 105]. For the purposes of this study, total *Bdnf* mRNA (exon IX) levels were evaluated due to differential hydroxymethylation analysis covering the whole gene body, and the variance by isoform specific expression was not accounted for. *Tlr4* signaling pathways have been shown to be increased with seizures [37, 55]. Loss of *Gad2* activity promotes seizures, *Gad2* knockout mice develop epilepsy, and humans with anti-GAD antibodies present with drug-resistant epilepsy [18, 51, 53, 100]. However, *Gad2* has also been reported to be increased in GABAergic neurons in epileptic rodents [25], showing the importance of cell type on gene expression. The inverse effect of 5-hmC on gene expression could be explained by the heterogeneity of cell types in the tissue, the contribution of other transcriptional regulators, or due to a regulatory effect of intergenic 5-hmC.

The role of intergenic 5-hmC in the regulation of epilepsy-related genes remains to be understood. In the present study, we observed that the majority of 5-hmC loss occurred in intergenic regions. Several mechanisms are described in the literature on how intergenic 5-hmC can affect gene expression depending on the genomic context. For example, enrichment of intergenic 5-hmC nearest to gene TSS has been shown to be associated with bidirectional regulation of gene expression [52, 121]. Several studies have reported 5-hmC association with H3K4me1 and enhancers, resulting in increased expression of the associated gene [17, 41, 58]. Additionally, intergenic 5-hmC has been shown to interact with CTCF sites, TADs, and transposable elements [14, 58]. Interestingly, DhMRs were associated with several epilepsy-associated GO terms, including GABA signaling regulation of ion transport, suggesting that 5-hmC maybe involved in regulating these pathways. Future research into the regulatory potential of 5-hmC in epilepsy is needed to fully understand the effect of 5-hmC on epilepsy permissive genes and what hippocampal cell types primarily are affected by 5-hmC.

To date, the postnatal role of TET enzymes in the healthy hippocampus and the consequence of aberrant TET enzyme activity and expression in epilepsy is poorly understood. In prior studies we investigated TET expression in TLE animals and found significant decreases in *Tet1* mRNA levels in area CA3 of the hippocampus and the dentate gyrus following status epilepticus, while 5-hmC is lost in area CA3 following status epilepticus and the dentate gyrus of TLE animals [105]. These findings suggest that TET1 and 5-hmC regulate neuronal activity differently across the timeline of epileptogenesis as well as hippocampal subfields. Recently, it has been reported that *Tet2* is increased in the brains of patients with drug-resistant epilepsy [59]. Further evidence for the importance of TET enzymes in epilepsy, humans with *TET3* mutations resulting in *TET3* deficiency can experience seizures [5, 69, 110].

We show that drug-resistant TLE patients have a loss of total 5-hmC in their hippocampi. Targeting 5-hmC directly or through modulation of TET activity could be a target for antiepileptic therapeutics in these patients. In addition, there are no drugs to date that treat the comorbidities of epilepsy such as memory loss. Previous studies have shown that Tet1 and DNA 5-hmC are crucial for learning and memory. *Tet1* isoforms differentially regulate fear memory [31], *Tet1* KO mice have deficits in fear memory extinction and LTD [104], *Tet1* overexpression impairs long-term memory formation [50]. *Tet2* overexpression in aged animals enhances fear memory [28], and in a mouse model of Alzheimer’s disease *Tet2* knockdown impairs cognition while *Tet2* overexpression improves memory [70]. In addition, global DNA 5-hmC is increased in area CA1 of the hippocampus following fear memory retrieval [125].

Our *in vivo Tet1* manipulations confirm that DNA hydroxymethylation changes can influence seizure vulnerability and resilience. We show that 5-hmC loss is sufficient to drive seizures and increasing 5-hmC promoted seizure resilience, thus the study of 5-hmC mediated mechanisms may provide novel insights into the hyperexcitable state of the epileptic brain. Targeting epigenetic pathways with pharmaceutical intervention is growing in interest for the treatment of several diseases including cancer [48, 120], and in epilepsy, recent studies have investigated the potential of histone deacetylase (HDAC) inhibitors in animal models of epilepsy with some success [16, 42, 114]. Targeting the TET/5-hmC pathway in epilepsy could potentially be a novel therapeutic that can treat both seizures and memory deficits. To further investigate the influence of DNA hydroxymethylation on hyperexcitability in epileptogenesis, and chronic epilepsy, future studies are needed. Finally, this study demonstrates that DNA 5-hmC should be considered when examining DNA methylation in epilepsy and provides additional insights that may help us to better interpret previous bisulfite methylation sequencing results.

## LIST OF ABBREVIATIONS

5-mC: 5-methylcytosine
5-hmC: 5-hydroxymethylcytosine
CA: Cornu ammonis
CGI: CpG island
CpG: C-phosphate-G-
CTCF: CCCTC-binding factor
DhM: Differentially hydroxymethylated
DhMRs: Differentially hydroxymethylated regions
DNMTs: DNA methyltransferase
DMRs: Differentially methylated regions
GABA: Gamma-aminobutyric acid
GO: Gene ontology
HDA: Histone deacetylase
KEGG: Kyoto encyclopedia of genes and genomes
KO: Knockout
LTD: Long-term depression
meDIP: Methylated DNA immunoprecipitation
meDIP-seq: Methylated DNA immunoprecipitation sequencing
scRNA-seq: Single cell RNA sequencing
TAD: Topologically associated domain
TET: Ten-eleven translocation
TLE: Temporal lobe epilepsy
TSS: Transcription start site

## DECLARATIONS

### Ethics approval and consent to participate

All animal procedures were approved by the University of Alabama at Birmingham’s institutional animal care and use committee (IACUC). Human tissue collection procedures were approved by the University of Alabama at Birmingham’s institutional review board (IRB).

### Availability of data and materials

The hMeDIP-seq data has been deposited to NCBI’s Gene Expression Omnibus (GEO) under accession number: GSE198737. The hMeDIP-seq analysis pipeline has been uploaded to GitHub.

### Authors’ contributions

R.H. wrote the manuscript. R.P., F.L., R.G.S. and L.V.H planned experiments. R.P., R.G.S., S.S.J., and

R.J.S. performed experiments. R.H., S.S.J., L.I, and R.B. generated figures. R.H., R.G.S., S.S.J., and L.I. analyzed data. All authors reviewed and edited manuscript.

## Acknowledgements

We would like to thank Drs. Kristen Riley, MD and Tore Eid, MD, of the Department of Neurosurgery at UAB, and the Department of Laboratory Medicine and of Neurosurgery at Yale School of Medicine respectively for their contributions in obtaining and sharing the human samples used for these studies. The bioinformatics analysis of this work was supported by the University of Alabama at Birmingham Biological Data Science Core, RRID:SCR_021766.

---

**Supplemental Figure 1.**

Altered Hydroxymethylation at both promoters and bodies of genes involved in neuronal and synaptic regulation.

a) Bar plot of selected genes with 5-hmC BAM counts changes at gene promoters and

b) Gene body. *p<0.05, **p<0.01, ***p<0.001 unpaired t-test.

**Supplemental Figure 2.**

Gene Pathways in association with Hydroxymethylation in Epilepsy

Heatmap shows molecular pathways influenced by differentially Hydroxymethylated genes of epileptic and non-epileptic rats with normalized 5-hmC BAM count.

a) Neurogenesis

b) Synaptic signaling

c) microRNA

d) Learning & memory

e) Genes regulated by nuclear calcium signaling.

**Supplemental Figure 3.**

Candidate Gene Descriptions

a) Schematic illustrating candidate gene selection. Significantly hypo- or hydroxymethylated genes with an adjusted p-value <0.05 were filtered for genes known to be associated with epilepsy or seizures. The epilepsy/seizure associated genes with the largest fold change were filtered by a Log2FC of at least 0.9 to generate a list of 26 genes candidate genes for regulation by 5-hmC in epilepsy.

b) Log2FC candidate genes in epilepsy relative to non-epileptic animals.

c) BAM counts of candidate genes in epileptic and non-epileptic rats.

d) Relative mean expression of candidate genes in astrocytes, glutamatergic neurons, gabaergic neurons, oligodendrocytes, and endothelial cells. Mean cell type expression values from the Allen Brain cell type database were collected and sorted into 3 bins for high, medium, or low average gene expression.

