## Supplementary figures and images for "Alterations in DNA 5-hydroxymethylation Patterns in the Hippocampus of an Experimental Model of Refractory Epilepsy"

### Supplemental Figure 1

Supplemental Figure 1

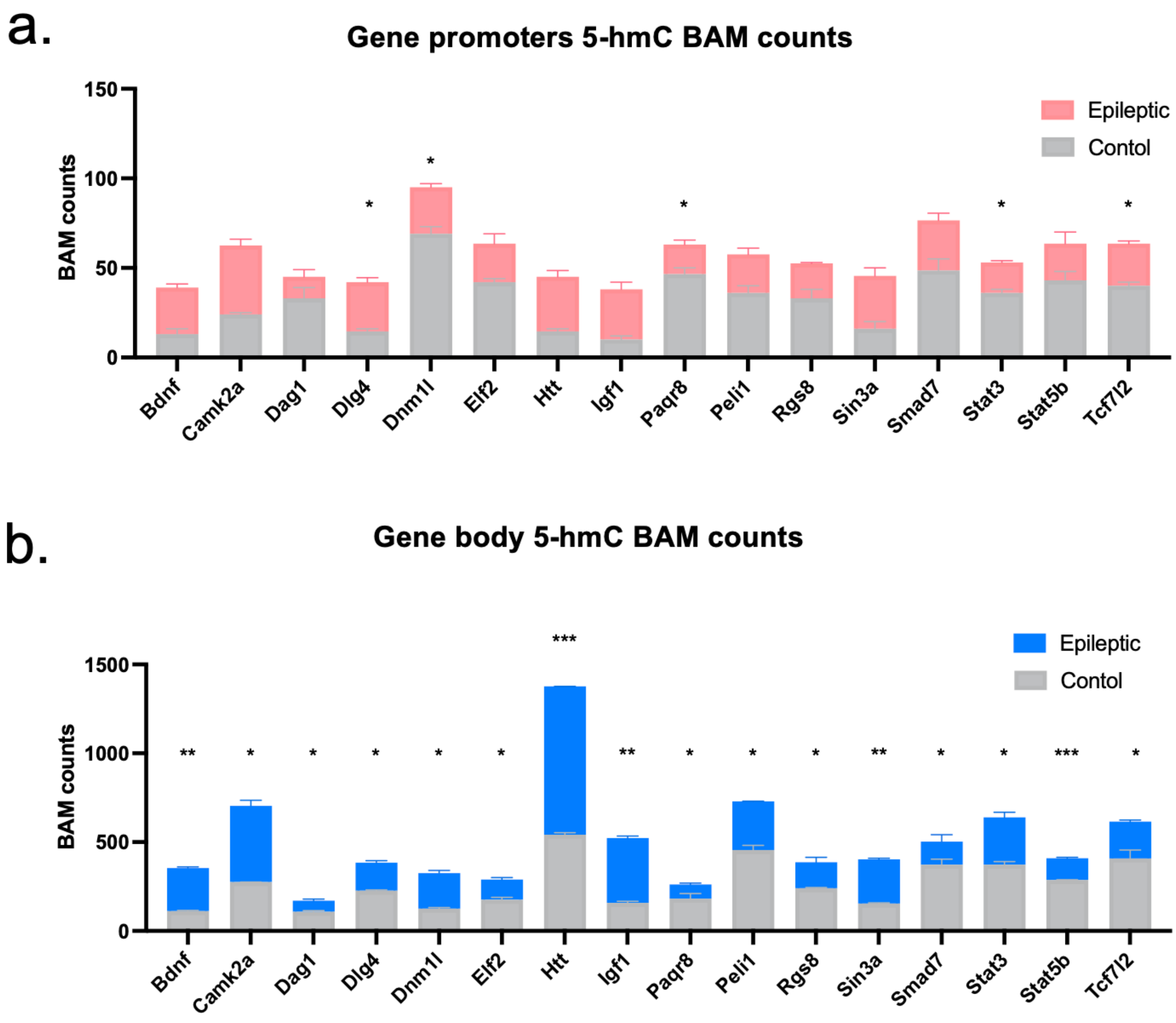

### Supplemental Figure 2

Supplemental Figure 2

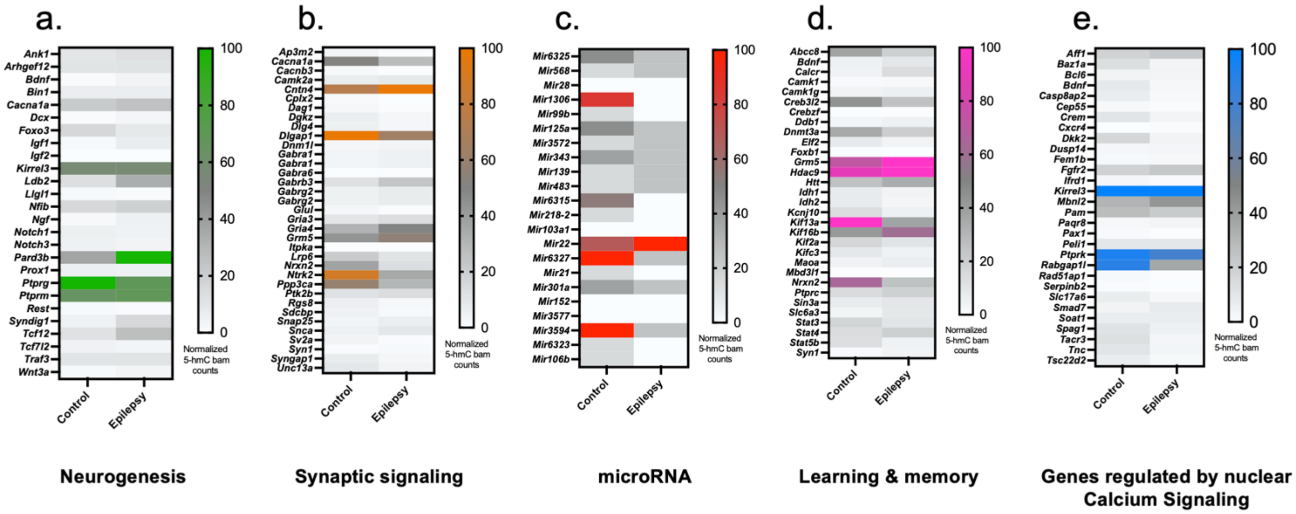

### Supplemental Figure 3

Supplemental Figure 3

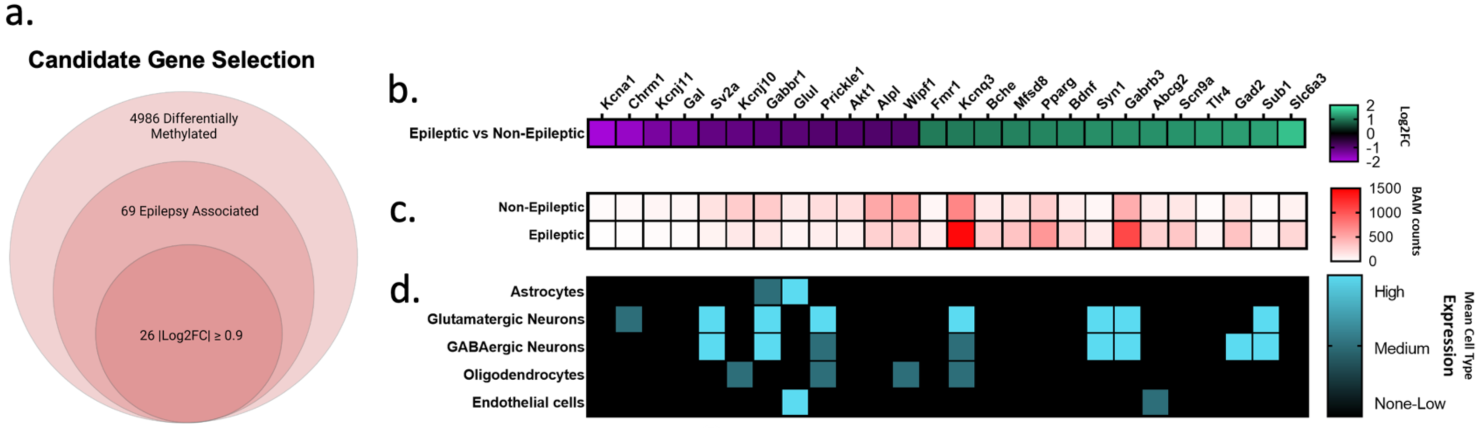
